# Interactions between genetic variation and cellular environment in skeletal muscle gene expression

**DOI:** 10.1101/105429

**Authors:** D. Leland Taylor, David A. Knowles, Laura J. Scott, Andrea H. Ramirez, Franceso Paolo Casale, Brooke N. Wolford, Li Guan, Arushi Varshney, Ricardo Oliveira Albanus, Stephen C.J. Parker, Narisu Narisu, Peter S. Chines, Michael R. Erdos, Ryan P. Welch, Leena Kinnunen, Jouko Saramies, Jouko Sundvall, Timo A. Lakka, Markku Laakso, Jaakko Tuomilehto, Heikki A. Koistinen, Oliver Stegle, Michael Boehnke, Ewan Birney, Francis S. Collins

## Abstract

From whole organisms to individual cells, responses to environmental conditions are influenced by genetic makeup, where the effect of genetic variation on a trait depends on the environmental context. RNA-sequencing quantifies gene expression as a molecular trait, and is capable of capturing both genetic and environmental effects. In this study, we explore opportunities of using allele-specific expression (ASE) to discover *cis* acting genotype-environment interactions (GxE) - genetic effects on gene expression that depend on an environmental condition. Treating 17 common, clinical traits as approximations of the cellular environment of 267 skeletal muscle biopsies, we identify 10 candidate interaction quantitative trait loci (iQTLs) across 6 traits (12 unique gene-environment trait pairs; 10% FDR per trait) including sex, systolic blood pressure, and low-density lipoprotein cholesterol. Although using ASE is in principle a promising approach to detect GxE effects, replication of such signals can be challenging as validation requires harmonization of environmental traits across cohorts and a sufficient sampling of heterozygotes for a transcribed SNP. Comprehensive discovery and replication will require large human transcriptome datasets, or the integration of multiple transcribed SNPs, coupled with standardized clinical phenotyping.

## Introduction

A substantial fraction of variability in gene expression is controlled by changes in transcription rates, mainly mediated by transcription factor (TF) proteins binding to specific DNA sequence motifs that define regulatory elements [1,2]. The abundance of such proteins and their regulatory co-factors may in turn be controlled by intrinsic mechanisms inherent to a cell, such as an individual's genetic makeup or regulatory programs specific to a cell type, as well as cellular responses to environmental cues. A regulatory element, defined by the DNA region recognized by a DNA-binding TF and other required transcriptional machinery, may be either intrinsic or environment-dependent. In intrinsic elements, the TF and binding machinery is controlled by cell-intrinsic mechanisms that operate within a closed system and are unresponsive to environment. By contrast, in environment-dependent elements the TF and binding machinery is responsive to an environmental stimulus. Both regulatory element types are susceptible to perturbation by genetic variation because the region recognized by the TF is encoded in the DNA sequence.

Many genetic studies document the effects of genetic perturbations of regulatory elements on gene expression - expression quantitative trait loci (eQTL) [3–6]. Although it is possible to detect *trans* (different physical chromosome) effects, eQTLs are typically identified within a local window, centered on the transcription start site (TSS), and assumed to act via *cis* (on the same physical chromosome) mechanisms. Variation in intrinsic regulatory programs is expected to give rise to such “standard eQTLs”, identified by modeling genetic effects on gene expression. However, it is also likely that variation in environment-dependent elements will be detected in standard eQTL studies, as it is unlikely for a variant to change the relationship between gene expression and environment without altering the mean gene expression levels for each genotype. Therefore we would expect a subset of eQTLs detected by modeling only genetic effects to also have effects unique to an environmental context. If one were to model the combined environmental and genetic effects on gene expression, such variants would exhibit interaction effects between genotype and environment (GxE) and could be described as GxE interaction quantitative trait loci (abbreviated as iQTLs in this paper), a specific type of eQTL whose effect changes according to an environmental context. To date, the overlap between standard eQTL and iQTL in human is largely unknown, as few studies have co-measured environmental and genetic effects at scale, and the technology for mapping such iQTLs is in its infancy.

In human populations, several GxE signals have been reported across diseases for various quantitative traits (reviewed in [7]), but few have mapped transcriptional iQTLs on a large scale, treating gene expression as a molecular quantitative trait [8–17]. Indeed transcriptional GxE effects have primarily been studied in model organisms where the environment and genotype can be controlled [18–23]. The challenge of mapping iQTLs using transcriptomic data outside of controlled laboratory settings lies in the confounding effects of environmental, biological, and technical factors on gene expression data, and the difficulty in isolating and/or accounting for such effects while preserving effects of the environment of interest.

However, such limitations may be mitigated if a study quantifies gene expression using RNA-seq technology because RNA-seq enables the measurement of allele specific expression (ASE), an alternative readout less prone to the confounders of gene level measurements [10, 24]. By quantifying differences in expression between haplotypes in samples heterozygous for a transcribed allele(abbreviated tSNP in this paper), ASE provides an internally controlled measurement where biological and technical exposures on the cells are essentially identical for both haplotypes. This makes ASE ideal for iQTL mapping since it minimizes batch effects while preserving *cis*-mediated environmental effects.

Furthermore, when integrated with standard gene expression data between individuals (abbreviated to gene-level expression in this paper), the two data types can serve as orthogonal forms of signal to validate iQTLs. In cases of true *cis* regulation of gene expression, when a TF preferentially binds to one allele, we would expect to observe increased ASE in participants heterozygous for the regulatory SNP. As an example, Fig 1 shows the different types of potential regulatory elements and the impact of different polymorphisms in schematic form. At the gene expression level, we would expect an iQTL to have different effects across environmental contexts in a genotype specific manner. In the ASE data, we would expect correlation between ASE and the environment only in individuals heterozygous for both the iQTL-SNP and tSNP. As opposed to standard eQTLs, which can be summarized by box-plots stratified by genotype, we believe a 6-panel regression plot is the most informative, and examples of expected behavior are shown in Fig S1.

**Fig 1.**
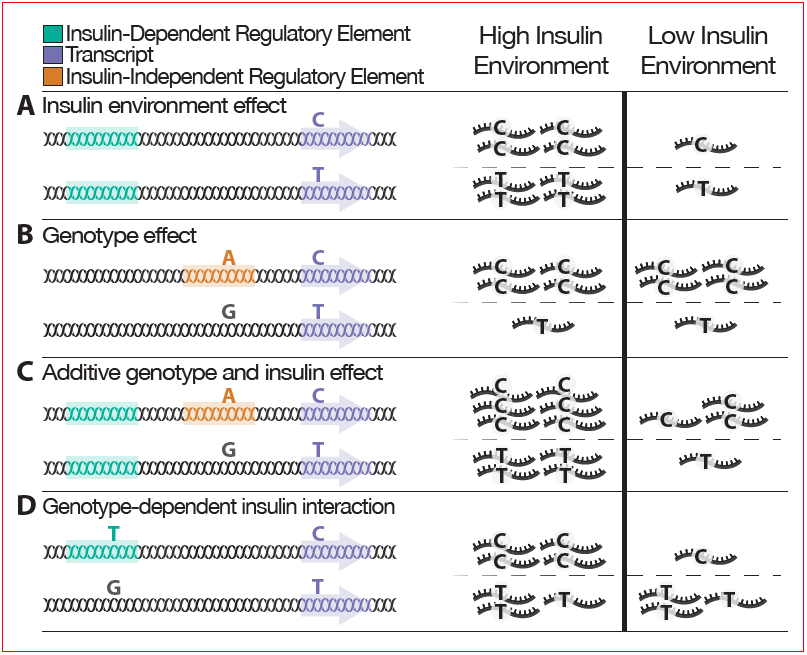
Genetic and environmental effects on gene expression. Blood insulin levels represent a cellular environment for tissues such as skeletal muscle. The left panel depicts a single genome with color-coded genomic elements and various heterozygous sites. The right panel shows the relative transcript abundance for the corresponding locus on the left panel. Some genomic elements contain genetic variants. When the variant is the same color as the element, the element is active. In some cases the variant is black, indicating that the variant renders the regulatory element nonfunctional and only basal transcription occurs. The purple element represents a gene with a transcribed SNP (tSNP), shown in the transcripts. Allele specific expression is calculated across both chromosomes and compared to the high and low environment.(A) When regulated by an insulin-responsive element (green), gene expression changes according to insulin concentrations in the extracellular environment. (B) When regulated by an insulin-independent element (orange) containing genetic variation, gene expression changes according to the presence of a genetic variant (eQTL), but not to insulin levels. The tSNP shows allelic bias due to the eQTL effect, but is not associated with the insulin environment. (C) When regulated by both an insulin-responsive element and an insulin-independent element containing genetic variation, the effects of the insulin environment and the genetic variation on gene expression may be additive, although more complex relationships are possible. The tSNP shows some imbalance due to the eQTL effect and is associated to insulin levels. Such cases may be identified as weak iQTLs. (D) When regulated by an insulin-responsive element containing genetic variation, there may exist an interaction effect between the genetic variant and insulin levels such that changes in gene expression across insulin environments depend on the genetic variant. The tSNP shows allelic imbalance associated with insulin levels due to the iQTL effect. One of several possible interaction effects depicted.

In this study, we explore the opportunities and challenges for iQTL mapping and replication using gene-level expression and ASE data. We illustrate our approach using RNA-seq from 267 skeletal muscle biopsies from the Finland-United States Investigation of NIDDM Genetics (FUSION) tissue biopsy study [25], as this dataset features RNA-seq co-measured with rich clinical phenotypes spanning blood metabolites, anthropometric measurements, and medication (S1 Table). Collectively, we treat all clinical phenotypes as “environmental traits” since we model skeletal muscle gene expression and therefore the response of a population of cells to the surrounding cellular environment - adjacent cells, extracellular matrix, blood plasma, and interstitial fluid - approximated by each phenotype.

As one clear limitation is sample size, we reduce the multiple testing burden by only testing eQTLs for GxE signals, based on the assumption outlined above that at least some of the strongest iQTLs will also show effects on mean gene expression when stratified by genotype and be detected also as eQTLs. With a well-calibrated statistical test, we identify 12 GxE signals that span 10 candidate iQTLs at a trait-specific FDR of 10%. Replication of such findings is challenging because of the lack of human studies on equivalent tissues with equivalent environmental measurements; however, two of the three testable traits shared with the larger GTEx study show non-random aggregate replication, although the need to restrict to heterozygous individuals limits the extent of this replication. This study highlights the utility of ASE based GxE analysis in observational studies, and emphasizes the need for large RNA-seq cohorts with standardized clinical phenotypes to enable study comparison and replication.

## Results and Discussion

### iQTL Results

As candidate iQTLs for each gene, we tested the most significant skeletal muscle eQTL per gene for the 19,455 autosomal, protein coding genes with at least one significant eQTL from our previous study of 267 Finnish muscle samples [25]. We tested for interaction of these SNP-gene pairs with 17 clinical phenotypes (S1 Table) by jointly modeling the impact of genotype effects on gene level expression and ASE levels (Methods). The resulting p-value distributions are well calibrated (S2 Fig), with the vast majority of tested SNPs consistent with the null distribution. Using a 10% FDR per trait, we identify 10 candidate iQTLs across 6 traits (12 unique gene-environment trait pairs) (Fig 2; Table 1; S2 Table). Of the clinical variables considered, sex is unique in that GxE sex signals could be due to environmental (for example, circulating sex hormones) or intrinsic, within cell, effects due to differences in gene expression from the sex chromosomes.

**Fig 2.**
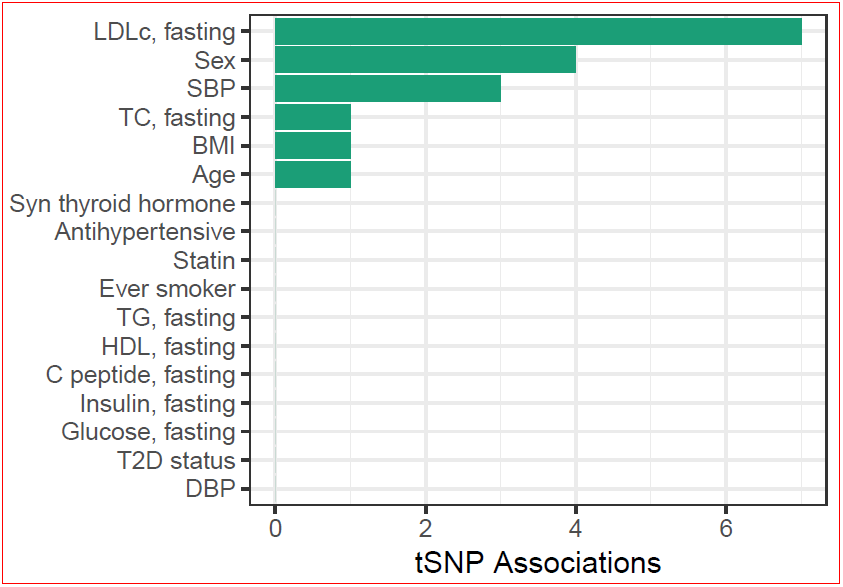
GxE signals. Number of tSNP-environment associations per clinical variable at a 10% FDR.

**Table 1.** IQTL results FDR 10%.

| Clinical Trait | Gene | Chr | tSNP position | iQTL alleles (ref/alt) | iQTL position | p-value ASE | p-value gene | p-value combined | q-value |
| --- | --- | --- | --- | --- | --- | --- | --- | --- | --- |
| Age | *PCNT* | 21 | 47786817 | G/T | 47823229 | 4.29x10<sup>-6</sup> | 1.25x10<sup>-1</sup> | 8.28x10<sup>-6</sup> | 0.0735 |
| Sex | *BSG* | 19 | 582775 | T/C | 572878 | 1.75x10<sup>-5</sup> | 1.00x10<sup>-1</sup> | 2.50x10<sup>-5</sup> | 0.0567 |
| Sex | *NRAP* | 10 | 115412793 | C/T | 115385650 | 1.65x10<sup>-7</sup> | 5.61x10<sup>-1</sup> | 1.59x10<sup>-6</sup> | 0.0136 |
| BMI | *DAGLB* | 7 | 6449272 | C/T | 6476915 | 3.54x10<sup>-2</sup> | 1.55x10<sup>-5</sup> | 8.48x10<sup>-6</sup> | 0.0753 |
| SBP | *ELP2* | 18 | 33750046 | T/G | 33743660 | 3.24x10<sup>-5</sup> | 3.58x10<sup>-2</sup> | 1.70x10<sup>-5</sup> | 0.0607 |
| SBP | *FHOD3* | 18 | 34324091 | T/C | 33970347 | 2.82x10<sup>-4</sup> | 5.07x10<sup>-3</sup> | 2.06x10<sup>-5</sup> | 0.0607 |
| SBP | *IGF2R* | 6 | 160453978 | T/C | 160379096 | 1.34x10<sup>-3</sup> | 9.18x10<sup>-4</sup> | 1.80x10<sup>-5</sup> | 0.0607 |
| TC, fasting | *AGMAT* | 1 | 15909850 | T/C | 15918676 | 2.52x10<sup>-3</sup> | 8.60x10<sup>-5</sup> | 3.54x10<sup>-6</sup> | 0.0315 |
| LDLc, fasting | *AGMAT* | 1 | 15909850 | T/C | 15918676 | 1.20x10<sup>-3</sup> | 4.82x10<sup>-4</sup> | 8.88x10<sup>-6</sup> | 0.0501 |
| LDLc, fasting | *DEPTOR* | 8 | 121061879 | G/T | 120930135 | 4.43x10<sup>-2</sup> | 1.69x10<sup>-5</sup> | 1.13x10<sup>-5</sup> | 0.0501 |
| LDLc, fasting | *FHOD3* | 18 | 34232657 | T/C | 33970347 | 6.78x10<sup>-3</sup> | 4.54x10<sup>-4</sup> | 4.21x10<sup>-5</sup> | 0.0623 |
| LDLc, fasting | *TMEM261* | 9 | 7799653 | A/G | 7830189 | 8.31x10<sup>-5</sup> | 1.39x10<sup>-2</sup> | 1.69x10<sup>-5</sup> | 0.0501 |

Summary of most significant tSNP for each iQTL-gene pair. Coordinates based on GRC37/hg19. The three p-value columns record the ASE, whole gene expressionlevel, and combined p-value respectively. The combined p-values are used for q-value calculation. Results with all iQTL-tSNP pairs are recorded in S2 Table.

### GTEx Replication

We sought to replicate these results using skeletal muscle data from the GTEx study http://www.gtexportal.org. Shared across studies, four traits were available for this purpose: age, sex, body mass index (BMI), and type 2 diabetes (T2D) status. Three of these variables: sex, BMI, and T2D status, had similar distributions in the GTEx and FUSION cohorts (S1 Table).

Despite significant differences in cohort populations, laboratory techniques, and analysis pipelines, we observe a trend in the replication rate of BMI and sex that increases with the significance of the iQTL in the FUSION discovery dataset (Fig 3). This trend was not observed in T2D, perhaps due to different criteria for inclusion of individuals with T2D. The FUSION tissue study only included individuals with newly diagnosed T2D, not yet treated with antihyperglycemic medications (described in [25]). In contrast, GTEx individuals may have had longstanding and heavily treated T2D [26, 27].

**Fig 3.**
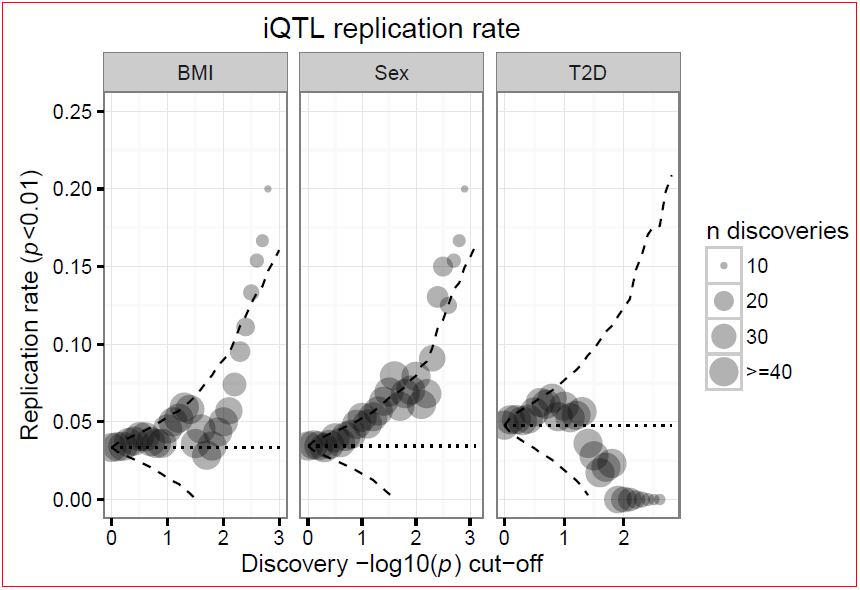
GTEx Replication. Replication rate (y axis) as a function of FUSION iQTL p-value cutoff (x axis). Dashed line represents two standard deviations from the null distribution, calculated using the hypergeometric distribution.

Although this bulk replication is reassuring, closer inspection of the BMI and sex trends revealed that two pairs of genes are driving the observed trend in both BMI and sex, highlighting the need of large sample sizes for such GxE analyses. To this point, only two significant iQTL-tSNP pairs from FUSION met the tSNP filtering criteria in GTEx (Methods), neither of which showed similar GxE effects, potentially indicating false positives (S3 Fig).

### Specific iQTL example: *FHOD3*

Despite the small number of reported hits and replication challenges, we observe some putative iQTLs with clear, consistent GxE effects in both gene expression and ASE data. The most clear, consistent example is *FHOD3*, formin homology 2 domain containing 3. *FHOD3* is essential for myofibril formation and repair, forming a doughnut shaped dimer, capable of moving along and extending actin filaments (reviewed in [28–30]). *FHOD3* is critical for heart development and function in mouse [31, 32] and fly [33] and exhibits tissue specific splicing patterns [34,35] shown to enable myofibril targeting in striated muscle [34,36].

We observed a GxE effect for *FHOD3* with both low-density lipoprotein cholesterol (LDLc) levels and systolic blood pressure (SBP) (Figure 4; S4 Fig). The LDLc association was discovered separately in the ASE of two tSNPs, spanning different exons (S Table 2; Figure 4; S4 Fig), while the SBP association was discovered with an additional tSNP, falling in an exon separate from the LDLc tSNPs. In addition, although not significant in the FUSION dataset, a GxE effect with BMI and *FHOD3* was one of the main drivers of the observed GTEx BMI replication trend (2.47x10^−4^ FUSION and 8.40x10^−4^ GTEx - minimum combined p-value across tSNPs). Evaluation of the raw data showed modest replication of the *FHOD3*-BMI signal between the FUSION and GTEx datasets (S5 Fig).

**Fig 4.**
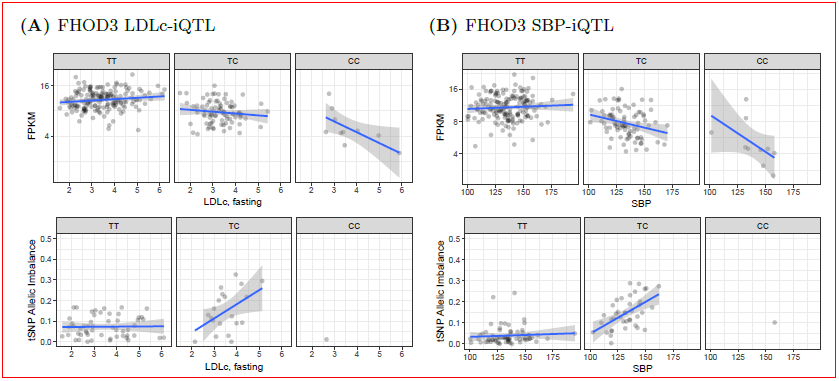
FHOD3 iQTL, rs17746240 (18:33970347). The data for each of the three possible iQTL genotypes are presented in separate plots (columns). The top row plots show the relationship between gene expression (y axis) and the clinical variable (x axis). The bottom row plots show the relationship between the allelic imbalance of the tSNP and the clinical variable (x axis). Note the bottom row has fewer samples because it is limited to samples heterozygous for the tSNP. (A) LDLc GxE effect with rs72895597 (18:34232657) as the tSNP (B) SBP GxE effect with rs2303510 (18:34324091) as the tSNP.

We previously calculated a muscle expression specificity index (mESI), comparing skeletal muscle expression to a reference panel of 16 diverse tissues, and binned these scores into deciles such that genes in the 1st decile are uniformly, lowly expressed and genes in the 10th decile are highly, specifically expressed in skeletal muscle [25]. We found *FHOD3* expression to be highly specific to skeletal muscle (mESI decile of 9). The iQTL tag SNP, rs17746240, and rs2037043, an additional SNP in high linkage disequilibrium (R^2^ = 0.99 in Finns from the GoT2D reference panel), overlap a skeletal muscle stretch enhancer (Fig 5A), a regulatory element shown to be a signature of tissue-specific active chromatin [37]. In addition, these variants fall in two distinct ATAC-seq peaks unique to skeletal muscle, an indicator of open chromatin (Fig 5B).

**Fig 5.**
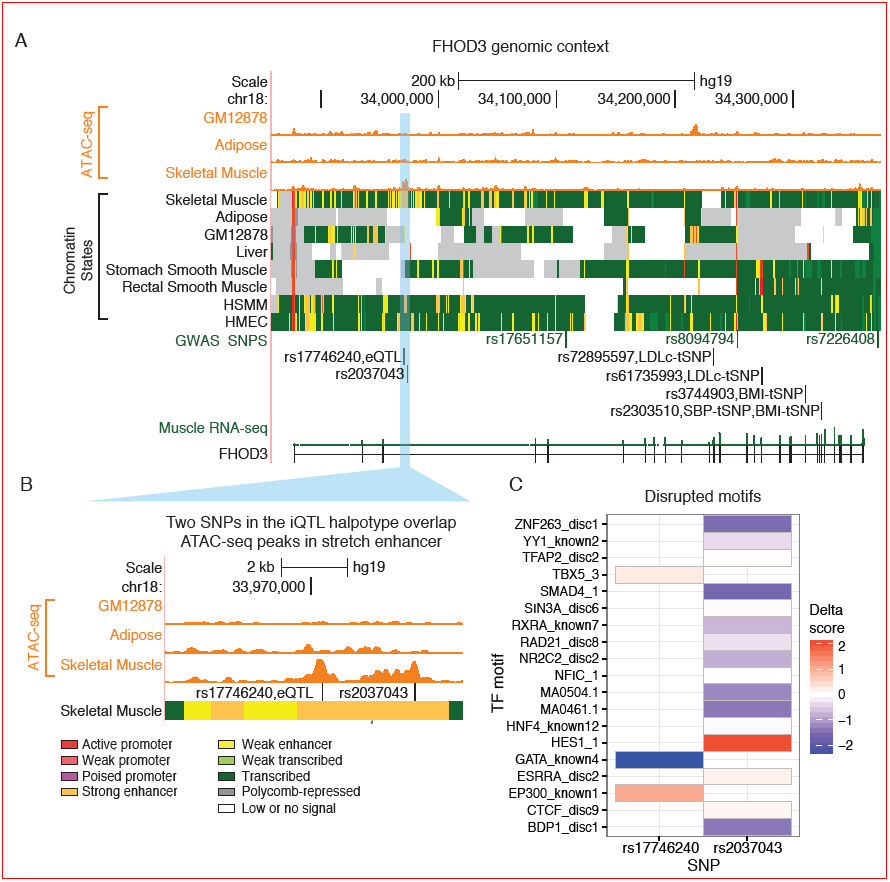
FHOD3 locus. (A) Top wiggle tracks show ATAC-seq signal in multiple cell types, followed by ChromHMM chromatin state tracks. Beneath are FHOD3 GWAS loci and the SNPs from this study (iQTL and tSNP). The bottom track shows the FUSION FHOD3 RNA-seq signal. (B) ATAC-seq signal highlights potential regulatory regions with the skeletal muscle stretch enhancer. (C) Effects of SNPs overlapping ATAC-seq peaks in the iQTL haplotype on in silico predicted TF binding.

Both SNPs affect predicted TF binding sites, as measured by the delta score (Methods). rs17746240 disrupts motifs for the GATA protein family, TBX5, and EP300 (Fig 5C). Within our skeletal muscle data, we find GATA2, GATAD1, GATAD2A, GATAD2B, and EP300 to be expressed (median FPKM > 1). The other variant, rs2037043, disrupts many motifs (Fig 5C) of which ZNF263, YY1AP1, YY1, SMAD4, SIN3A, RXRA, RAD21, NR2C2AP, NR2C2, NFIC, HES1, ESRRA, CTCF, and BDP1 are expressed in skeletal muscle (median FPKM > 1), making it difficult to identify a specific TF.

## Conclusion

Understanding the genetic regulators of molecular responses to environment, both at the cellular and organismal level, is essential for a complete understanding of the relationship between genotype and phenotype. Environmental influences are a critical part of human disease etiology, but are far harder to study than intrinsic genetic factors. RNA-seq technology provides an information-dense molecular readout that includes ASE, an internally controlled experiment that minimizes technical artifacts by comparing read counts *within* samples instead of *between* samples [10,24]. Because ASE reduces confounding effects present in gene-level data that are difficult to distinguish from environmental effects, ASE is an ideal molecular readout for probing GxE effects. This study, which is amongst the first to leverage ASE in humans to map GxE effects [10,13], demonstrates both the potential and the limitations for using ASE to unravel complex gene-environment regulatory structures. Using a well-calibrated model, we find a handful of iQTLs and show some level of bulk replication. Despite the low level of discovery in this study, which we believe is primarily limited by sample size, our success suggests that at least some eQTLs are likely to be in fact iQTLs.

This study highlights several challenges associated with using ASE signal for mapping regulatory loci. Such analyses require sufficient sampling of double heterozygotes of the iQTL and tSNP, and therefore large sample sizes are required for a well-powered study. Another limitation of ASE is that it can only be used to identify *cis*-effects. Previous studies indicate that many iQTLs operate distally, in *trans*, on highly regulated genes with more opportunities in the regulatory chain for genetic perturbation [8,15,22,23]. Because our method requires ASE, we could only assay local, *cis*-effects, and therefore may miss many large *trans*-effects.

In the future, we will need larger studies of specific human tissues with co-measured genetic, molecular, and clinical information. The possibility of mapping iQTLs underscores the importance of detailed characterization of study participants, especially when integrating molecular and genetic data with detailed clinical information. This becomes particularly relevant for replication studies, and argues for the standardization of a core set of phenotypes and environmental exposures between large cohorts. In addition, further development of statistical models to boost power will be needed - for instance by simultaneously modeling total gene expression and ASE, as well as accommodating technology developments, such as the integration of perfectly phased tSNP allele counts within a gene, made possible by long reads.

## Materials and Methods

Sample recruitment, muscle biopsy procedures, genotype processing, and RNA sequencing have been previously described [25].

### Ethics Statement

The study was approved by the coordinating ethics committee of the Hospital District of Helsinki and Uusimaa. A written informed consent was obtained from all the subjects.

### Phenotype Processing

Metabolites were measured after a 12-hour overnight fast, during a 4-point (0, 30, 60, 120 min) oral glucose tolerance test (OGTT) [25]. Serum triglycerides, total and HDL cholesterol were measured by enzymatic methods with Abbott Architect analyzer (Abbott Laboratories, Abbott Park, IL, USA). LDL cholesterol concentration was calculated using the Friedewald formula [38]. Serum insulin and serum C-peptide concentrations were assayed by chemiluminescent microparticle immunoassays using Architect analyzer. Patient medications were also recorded at time of OGTT. Patient medications were analyzed and categorized by physician review. All phenotypes considered are listed in S Table1.

We inverse normalized all continuous traits. Blood pressure measurements were missing from 2 participants, whose samples were dropped when analyzing blood pressure traits. Prior to fitting models, we regressed all continuous traits on age, age^2^, and sex, except for age where we regressed only on sex.

### ASE Processing

We quantified ASE in autosomal, protein coding genes as described previously [25], removing tSNPs that showed mapping bias based on simulated reads. To obtain a high confidence ASE dataset, we removed tSNPs per sample with < 30 total reads. We subsequently required that tSNPs were heterozygous in >= 20 samples. From the remaining 25,913 autosomal tSNPs, we discarded 1,254 tSNPs where one or more sample exhibited near mono-allelic expression, defined as | 0.5 - ( count ^alternate SNP^ / count ^total^) | > 0.4. Altogether, we considered 24,659 tSNPs to map candidate iQTLs.

### iQTL Discovery

Using 19,455 autosomal, protein coding skeletal muscle eQTLs published in [25], we tested for GxE effects in the ASE and gene expression data across all clinical traits. For ASE data, we used EAGLE [10], which models count overdispersion using a random effect term with per tSNP variance vs with an inverse gamma prior *IG(a, b)*. We learned the hyperparameters *a*, *b* for this distribution across all tSNPs after filters, estimating them to be 1.80, 0.0024 respectively. For sample *i* and tSNP *s*, we mapped GxE signals by fitting the model:

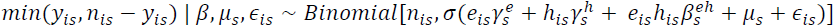

Here *n*
_
*is*
_ and *y*
_
*is*
_ denote the total and alternative read count for individual *i* at tSNP *s*, *e*
_
*is*
_ the environment, *h*
_
*is*
_ the indicator that the eQTL is heterozygous, *µ*
_
*s*
_ an intercept term to take into account unexplained allelic imbalance unrelated to the environment,

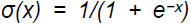

the logistic function, *ε*
_
*is*
_
*|v∼N(0, v*
_
*s*
_
) a per individual per locus random effect modeling over dispersion, and, 
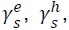
 and 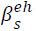
 the effect sizes of the environment, eQTL heterozygosity status, and SNP*environment interaction, respectively. We test the null hypothesis 
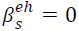
 using a likelihood ratio test. As covariates, we included the first two principal components (PCs) calculated across all genotypes, consistent with Scott *et al.* [25]. In our analyses we required ≥ 15homozygous and ≥ 15 heterozygous samples for the eQTL tag SNP and, in the case of dichotomous variables, no group was formed with < 5 samples. With these filters, we could only test for iQTL effects in a subset of genes that differed according to clinical trait in the case of discrete variables where the total sample size was not constant due to missing data (S6 Fig).

We also mapped GxE interaction effects for each candidate iQTL in total gene expression data using a linear model for expression levels, testing interactions for each gene-environment pair. Let y_j_ be a vector of inverse normalized FPKMs for gene *j* across individuals. We consider the following linear genetic model of gene expression:

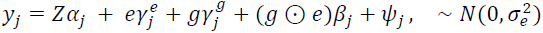

Here *Z* denotes the matrix design of fixed effect confounding covariates, *e* and *g* the environment and genotype vector, *g* ⊙ *e* their element-wise product, ψ_*j*_ Gaussian noise, and 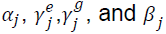 the effects of covariates, environment, genotype, and the genotype*environment interaction respectively.

To capture hidden variation in gene expression data, we used PEER [39, 40] as described previously [25] to learn latent factors. For covariates in the GxE interaction model, we included sequencing batch, the first two genotype PCs, and the first two PEER factors, as a recent report suggests two PEER factors capture the majority of technical variation, preserving biological effects [41]. We additionally include age and sex as covariates when either trait was not considered as an environmental trait. We implemented the GxE model using the linear mixed model framework LIMIX (v0.7.6) [42, 43].

We combined the ASE p-values and gene expression p-values using Fisher’s combined test. We controlled for FDR per environment using the Benjamini– Hochberg procedure [44]. Our method assumes 1) ASE and gene expression are independent measurements for GxE and 2) we have enough double heterozygous individuals to map the iQTL.

### GTEx Replication

We conducted a replication study using genotype, gene expression, and ASE from the GTEx v6 dbGaP release (phs000424.v6.p1). ASE was calculated across imputed genotypes of 360 skeletal muscle samples. The GTEx samples were collected post-mortem and do not have available many of the traits assayed in the FUSION samples. Of the clinical variables measured in the FUSION dataset, four were also recorded in the GTEx dataset - age, sex, BMI, and T2D status - from which we excluded age as the distribution was significantly different between FUSION and GTEx (S Table 1).

Notably, besides the differences in collected phenotype information and age distribution, the GTEx data differ from the FUSION data in four other relevant ways: 1) FUSION is drawn from a more genetically homogenous population (Finland); 2) FUSION is sequenced to mean depth of 91.3M reads per sample compared to 82.1M reads per sample in GTEx; 3) FUSION uses a 100bp strand specific, paired-end read protocol for RNA-seq and GTEx uses 76bp non-strand specific, paired-end RNA-seq; and 4) the computational analysis pipelines are different for read mapping, expression abundance quantification, and ASE calculations [45].

Within the GTEx dataset, we tested for GxE effects with the FUSION eQTL SNPs, using the ASE interaction and gene expression interaction models described above. Because our goal was replication of the FUSION genotype-environment interactions we did not require the eQTL to be significant. For the GTEx ASE interaction model, we including the first three genotype PCs as covariates, as was used previously by the GTEx consortium [45], and for the gene expression interaction model, we included age, sex, expression batch, the first three genotype PCs, and the first two PEER factors from the GTEx data release as covariates. We tested iQTL-tSNP pairs in GTEx with sufficient double heterozygotes to pass the filters described above. For genes with multiple tSNPs, we selected the minimum iQTL p-value per gene for the GTEx and FUSION datasets separately. Treating the FUSION data as a discovery dataset, we calculated the replication rate across varying p-value threshold cutoffs. We selected *n* FUSION hits at a given p-value cutoff from *N* total shared iQTLs without replacement, stopping when *n* < 10. At each cutoff, we calculated *k*, the number of FUSION hits that replicate in GTEx (GTEx p-value < 0.01), out of the total number of nominally significant GTEx hits, *K*. Using the mean, *K/N*, and the hypergeometric distribution, we estimated two standard deviations from the null distribution. Because we select the minimum iQTL-tSNP pair per gene it is possible that genes with more tSNPs will be more likely to show significant results. We calculated the average tSNPs for the replicated and not replicated iQTL sets to explore if sampling from a larger number of transcribed SNPs was responsible for the observed trends (S7 Fig).

**Figs7.**
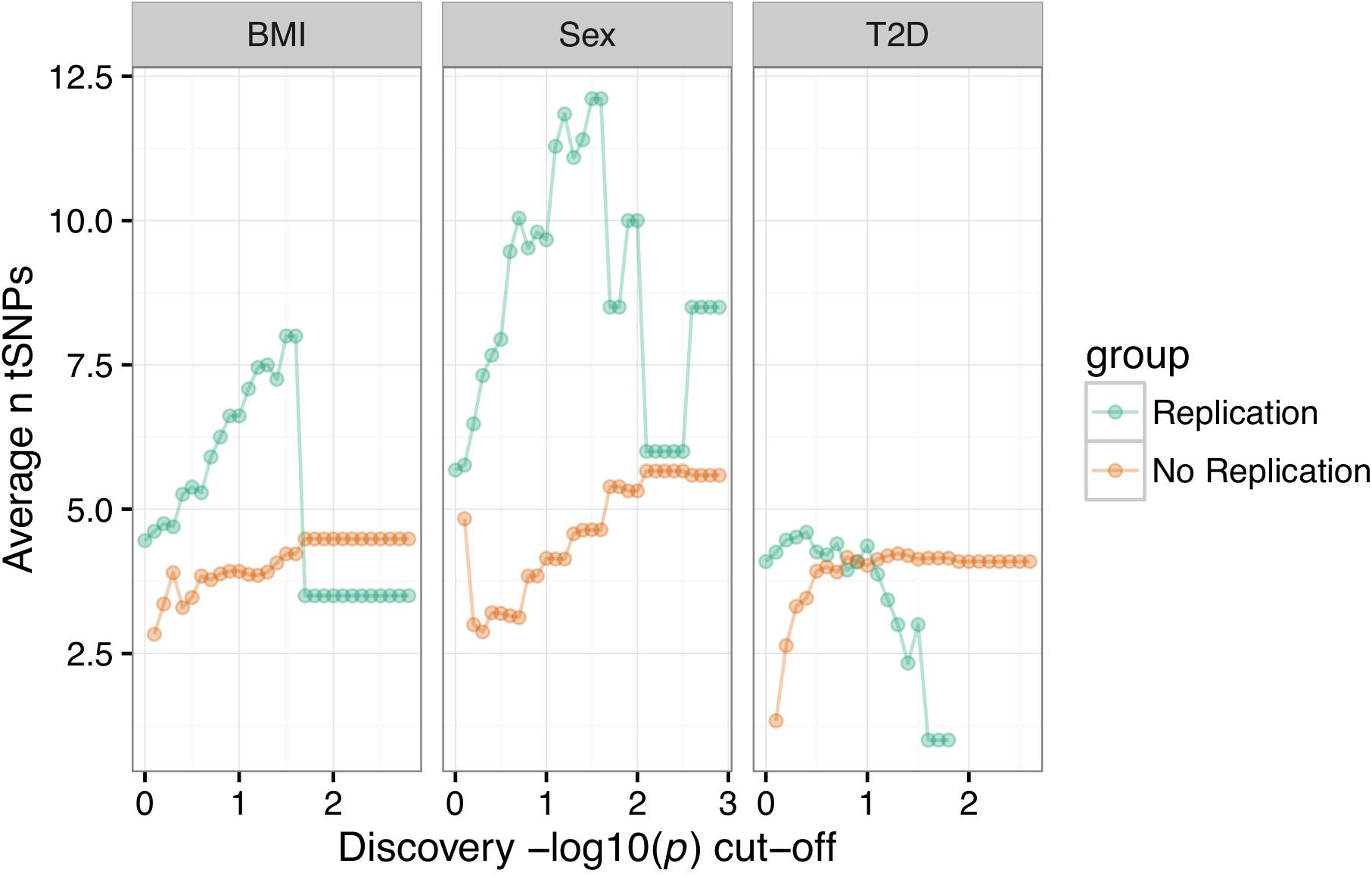
S7-GTEx Replication. Average number of tSNPs in the genes with signals that replicated (Replication group) and signals that did not replicate (No Replication).

### Chromatin States

We performed integrative chromatin state analyses as reported previously [37]. Briefly, we collected cell/tissue ChIP-seq (chromatin immunoprecipitation followed by sequencing) reads from a diverse set of publicly available data. Chromatin states were learned jointly by applying the ChromHMM (v1.10) algorithm at 200bp resolution to six data tracks (Input, K27ac, K27me3, K36me3, K4me1, K4me3) from each of the cell/tissue types [46,47]. We elected a 13 state model as it provided sufficient resolution to identify biologically meaningful patterns in a reproducible way [37].

### ATAC-seq Footprinting

Assay for transposase-accessible chromatin (ATAC-seq) generates detailed maps of open, active chromatin and TF binding dynamics [48]. We used previously published ATAC-seq data in skeletal muscle [25].

### Transcription Factor Binding Predictions

To identify potential transcription factor binding sites (TFBS), with particular attention to those that may be affected by variants, we generated short sequence fragments around each of the biallelic SNPs and short indels discovered in 1000 Genomes Phase 3 (release 5), by embedding each allele in flanking sequence (29bp on each side) from the hg19 human reference genome. We scanned the entire reference sequence, as well as these variant fragments, with a library of position weight matrices (PWMs) compiled from JASPAR [49], ENCODE [50], and Jolma *et al.* [51], using FIMO [52] from the MEME suite [53]. FIMO was executed using the background nucleotide frequency of the human reference (40.9% GC) and the default p-value cutoff, 10^−4^.

To quantify the effect of SNPs on these motifs, we calculated a delta score, -log10(p^alternate allele^)- -log10(p^reference allele^), for each SNP where at least one of the alleles passed our p-value cutoff of 10^−4^. In cases where a PWM hit was not detected for the second allele by FIMO at a threshold of 0.01, we use a value of 0.01 for that allele, so that the delta score will be conservative in these cases.

## Acknowledgements

We thank Anthony Kirilusha, John Didion, Daniel Bar, and Lori Bonnycastle for helpful comments and feedback. We also thank Julia Fekecs for help in designing Fig 5.

## Supporting Information

**S1 Table. Clinical traits.** Phenotype information used as traits from the FUSION tissue biopsy study participants and GTEx skeletal muscle participants. For T2D status in GTEx, only T2D status available, non-T2D participants presumed to be NGT. In some cases, the GTEx T2D status was missing (NA), therefore T2D fraction calculated over non-missing data.

**S2 Table. All iQTL-tSNP pairs FDR 10%.** All candidate iQTLs (FDR 10%).

**S1 Fig.**
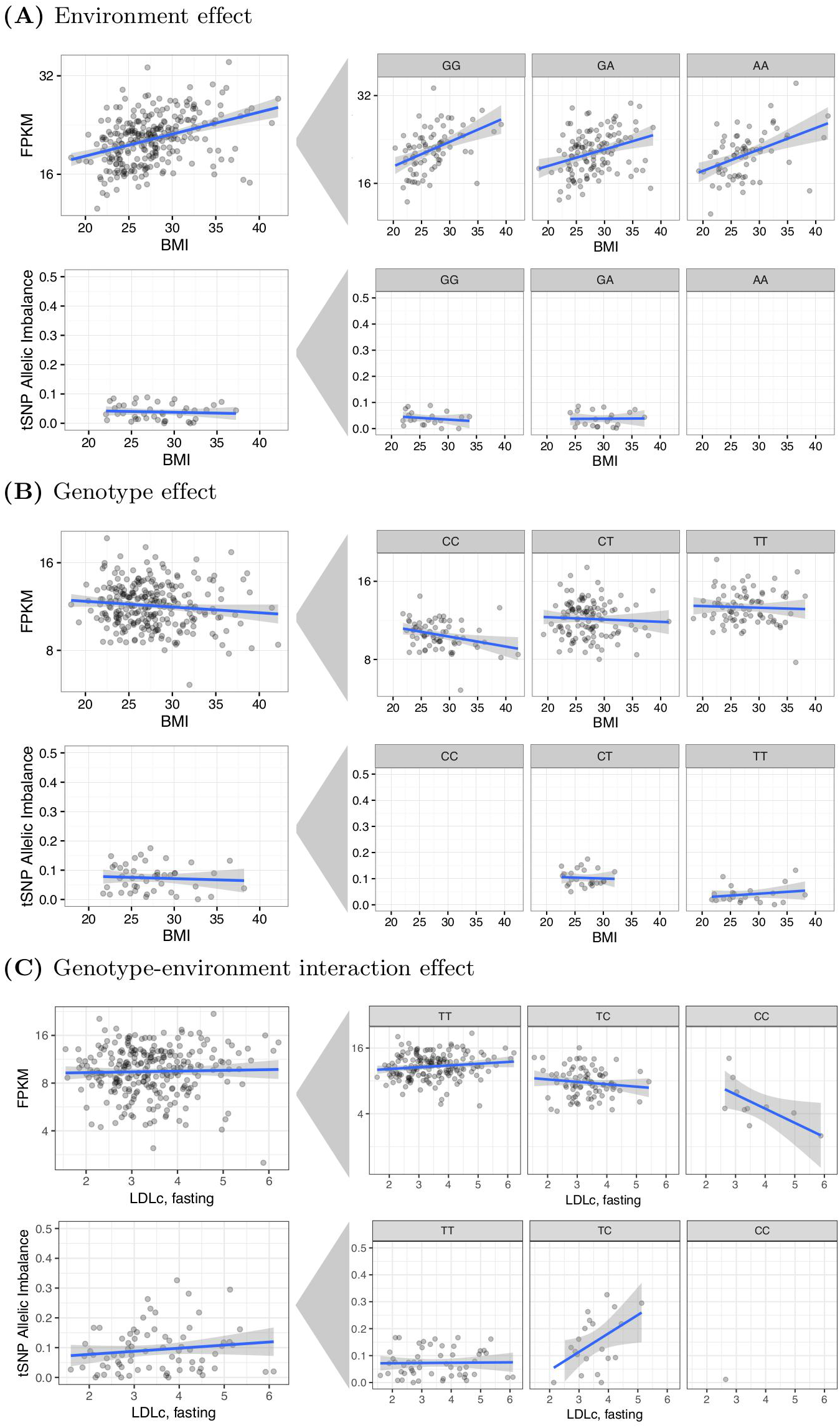
Examples of genetic and environmental effects. (A) Example of a pure environment effect in SZRD1 - rs12568938 regulatory SNP (rSNP) and rs7529767 transcribed SNP (tSNP). SZRD1 expression is associated with BMI, and the rSNP does not affect gene expression. The relationship between SZRD1 and BMI does not change across the rSNP alleles, and BMI is not associated with allelic imbalance. (B) Example of a pure genetic effect in RBM6 - rs9881008 regulatory locus and rs2023953 tSNP. BMI is not associated with RBM6 expression or allelic imbalance. The rSNP alleles are associated with RBM6 expression and allelic imbalance is increased in samples heterozygous for the rSNP. (C) Example of a GxE effect in FHOD3 - rs17746240 regulatory locus and rs72895597 tSNP. The relationship between LDLc and FHOD3 expression changes according to the rSNP allele as well as the overall expression abundance levels. LDLc is only associated with allelic imbalance in heterozygous individuals, where preferential TF binding could occur.

**S2 Fig.**
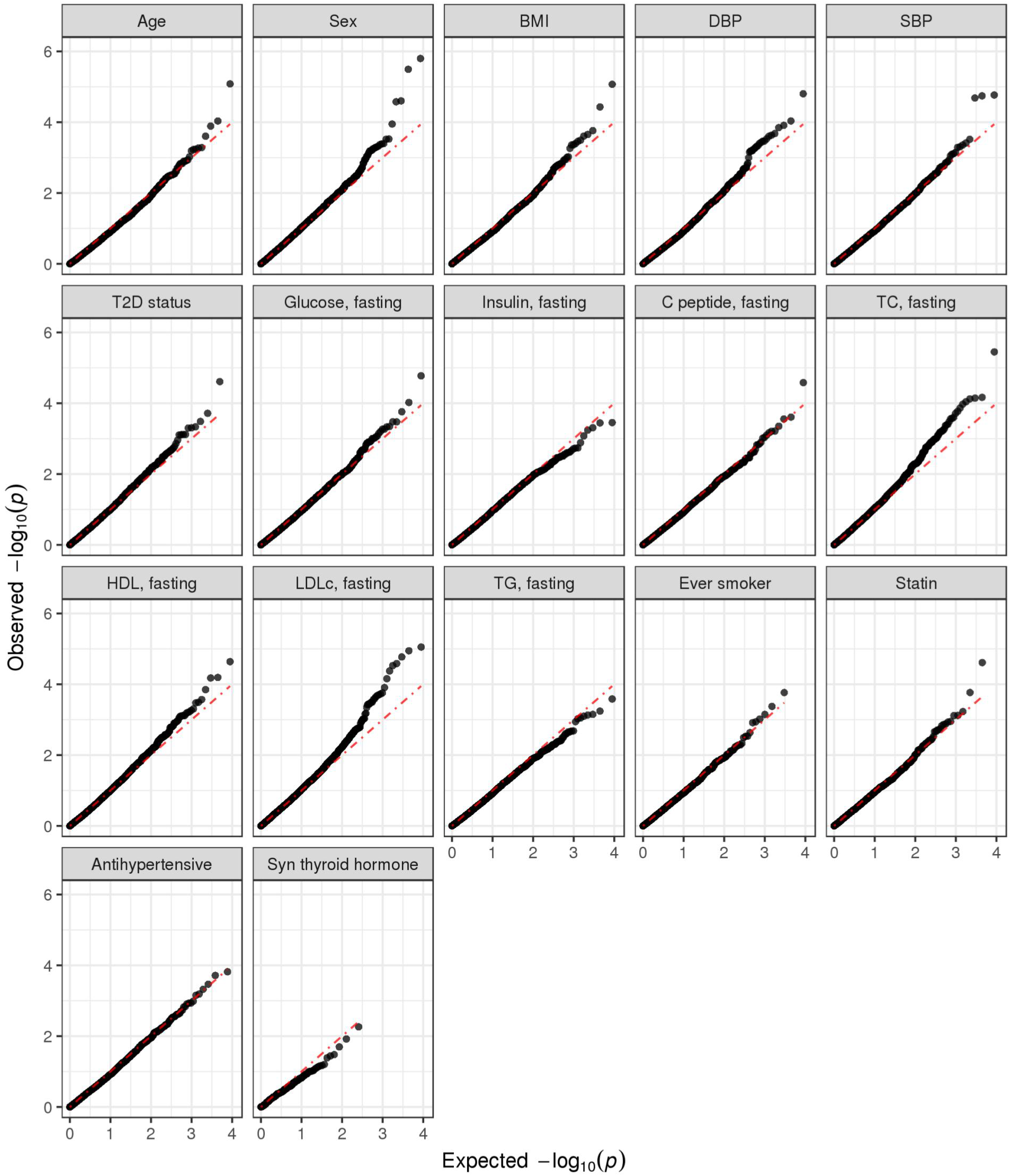
QQ-plots across traits. QQ-plots of GxE signal discovery across clinical traits.

**S3 Fig.**
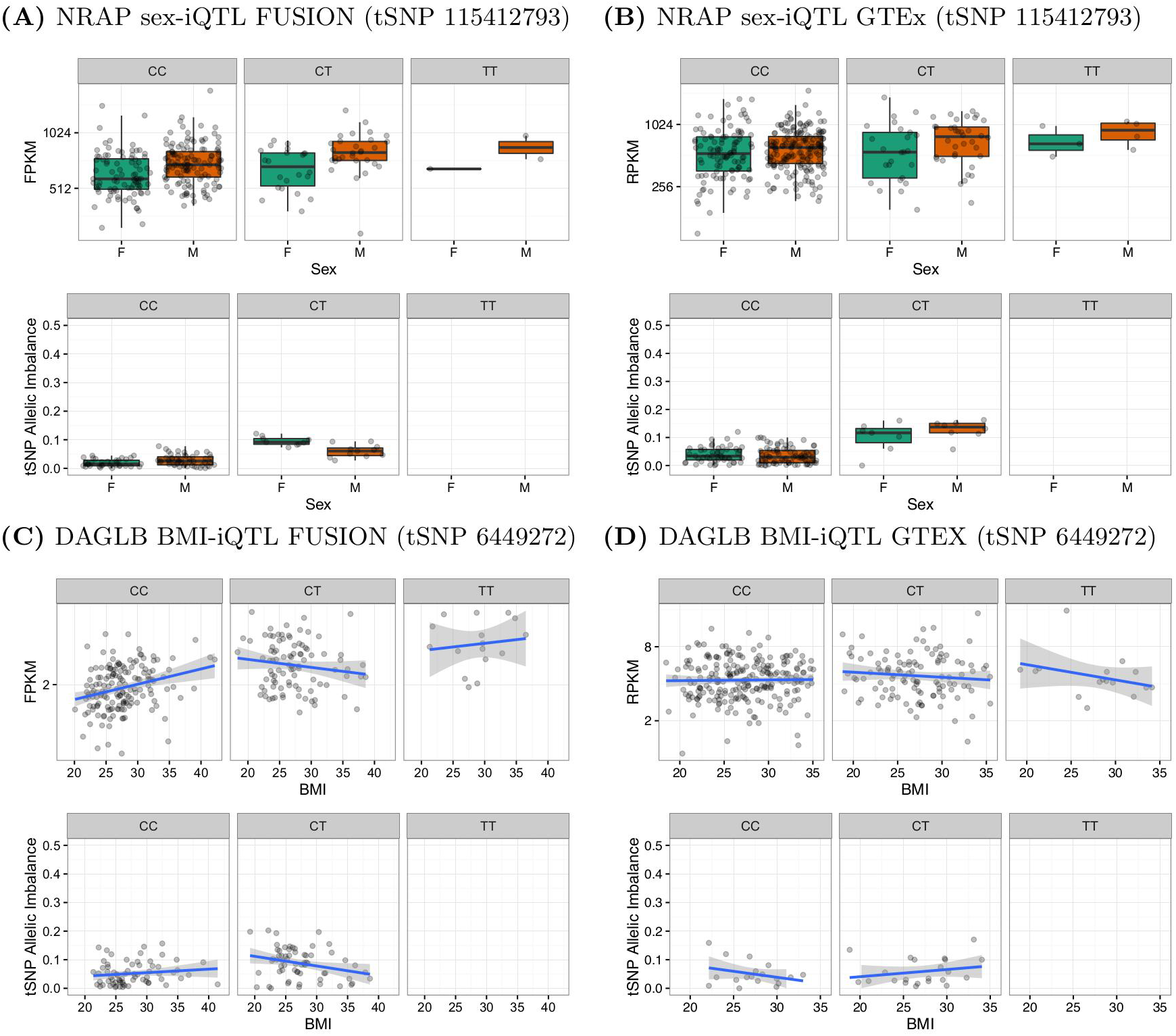
Comparison of candidate FUSION iQTLs to GTEx. (A) NRAP sex-iQTL in FUSION (B) NRAP sex-iQTL in GTEx (C) DAGLB BMI-iQTL in FUSION (D) DAGLB BMI-iQTL in GTEx.

**S4 Fig.**
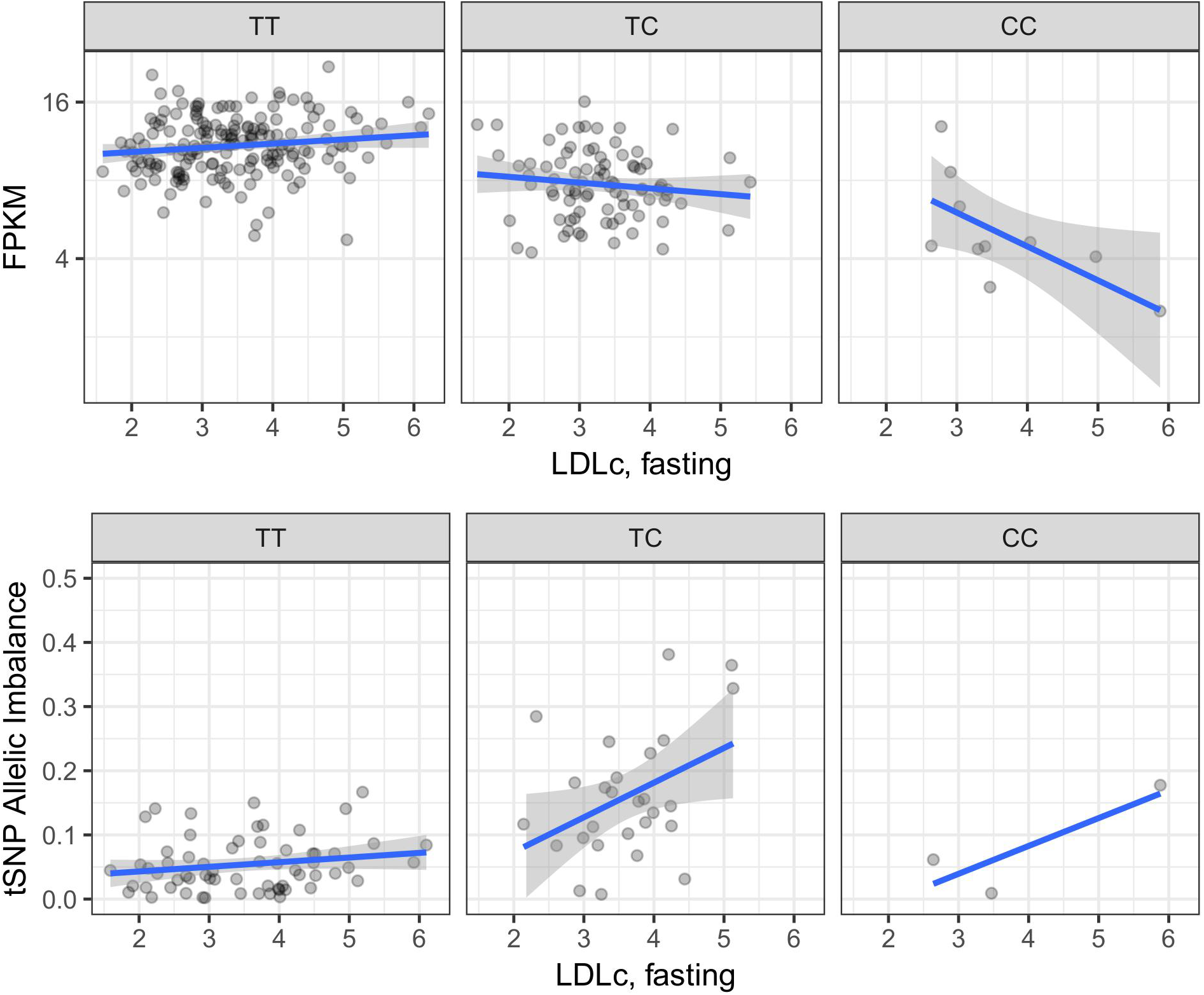
Additional FHOD3 LDLc-iQTL. Additional LDLc GxE effect with rs61735993 (18:34273279) as the tSNP.

**S5 Fig.**
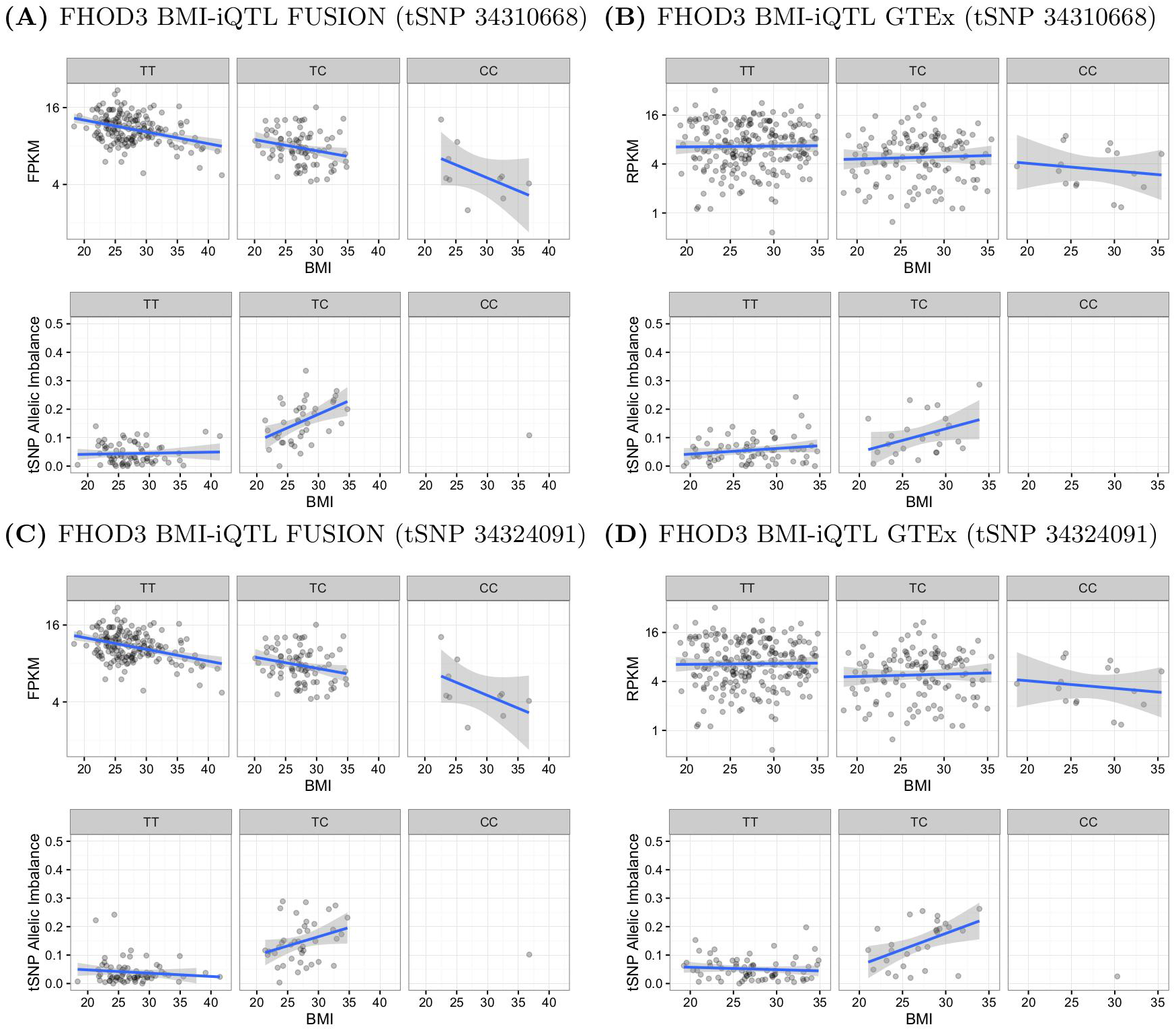
Comparison of FHOD3 BMI-iQTL in FUSION and GTEx. (A) FHOD3 BMI-iQTL in FUSION with rs3744903 (18:34310668) as the tSNP (B) FHOD3 BMI-iQTL in GTEx with rs3744903 (18:34310668) as the tSNP (C) FHOD3 BMI-iQTL in FUSION with rs2303510 (18:34324091) as the tSNP (D) FHOD3 BMI-iQTL in GTEx with rs2303510 (18:34324091) as the tSNP.

**S6 Fig.**
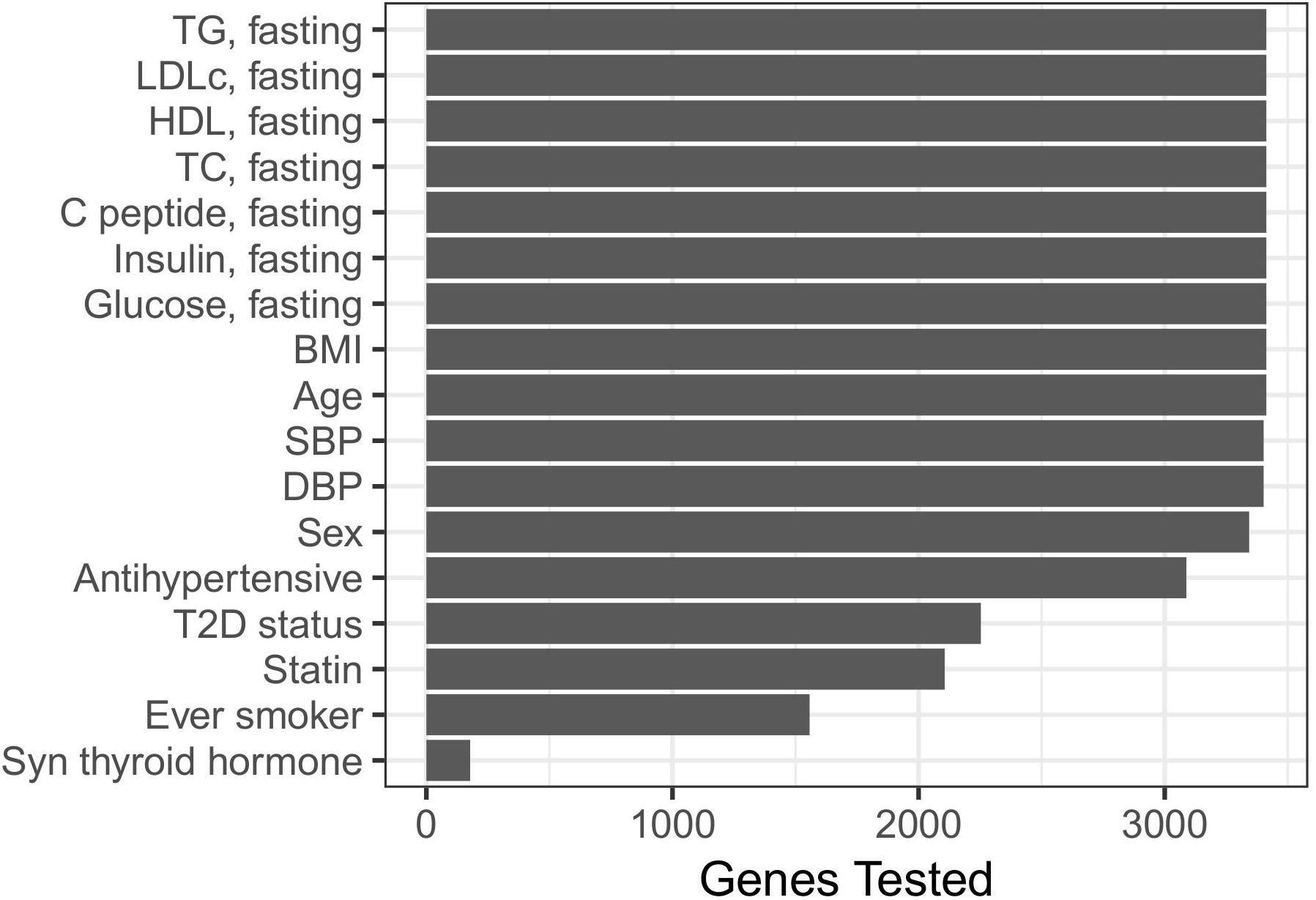
Total number of tested genes across traits. Total number of genes in FUSION considered for each clinical trait.

